# Regulation of eIF2α Phosphorylation by MAPKs Influences Polysome Stability and Protein Translation

**DOI:** 10.1101/2021.08.30.458160

**Authors:** Sana Parveen, Haripriya Parthasarathy, Dhiviya Vedagiri, Divya Gupta, Hitha Gopalan Nair, Krishnan Harinivas Harshan

**Author notes:** Equal Contributors (Sana Parveen, Haripriya Parthasarathy).

## Abstract

Regulation of protein translation occurs primarily at the level of initiation and is mediated by multiple signaling pathways, majorly mechanistic target of rapamycin complex 1 (mTORC1), mitogen-activated protein kinases (MAPKs), and the eukaryotic translation initiation factor eIF2. While mTORC1 and eIF2α influence the polysome stability, MAPKs influence the phosphorylation of the cap-binding protein eIF4E and are known to influence translation of only a small set of mRNAs. Here, we demonstrate that p38 MAPK and ERK1/2 regulate translation through integrated stress response (ISR) pathways. Dual inhibition (dual-Mi) of p38 MAPK and ERK1/2 caused substantial phosphorylation of eIF2α in a synergistic manner, resulting in near-absolute collapse of polysomes. This regulation was independent of Mnk1/2, a well-studied mediator of translation regulation by the MAPKs. Dual-Mi-induced polysome dissociation was far more striking than that caused by sodium arsenite, a strong inducer of ISR. Interestingly, induction of ISR caused increased p38 phosphorylation, and its inhibition resulted in stronger polysome dissociation, indicating the importance of p38 in the translation activities. Thus, our studies demonstrate a major, unidentified role for ERK1/2 and more particularly p38 MAPK in the maintenance of homeostasis of polysome association and translation activities.

## INTRODUCTION

Protein translation is a major regulatory event in gene expression in eukaryotes. Initiation is often the target for translation regulation and is achieved either by limiting the cap complex assembly or through preventing the recycling of pre-initiation complex (PIC) (1). Regulation is achieved through three major signal pathways, viz. mechanistic target of rapamycin complex 1 (mTORC1), initiation factor eIF2 and MAPK-eIF4E pathways. While alterations in the activities of mTORC1 and eIF2 directly impact polysome assembly, those in the eukaryotic translation initiation factor (eIF4E) do not appear to be globally consequential. eIF4E, the cap binding protein associates with eIF4G and eIF4A to form eIF4F complex whose formation is influenced by eIF4E availability (2). eIF4E is phosphorylated at S209 by MAPK interacting kinases Mnk1 and Mnk2 (3,4) that are regulated by two mitogen activated protein kinases (MAPKs), viz. extracellular signal regulated kinases (ERK1/2) and p38 MAPK (hereafter referred to as p38) (5,6). Even as the role of phosphorylation on the affinity of eIF4E for the cap is contradicted (7), phosphorylated eIF4E has transforming and oncogenic potentials (8–10). eIF4E overexpression is reported in large cases of cancers (11). It is proposed that eIF4F complex with phosphorylated eIF4E can translate certain transcripts that are excluded by the complex with unphosphorylated eIF4E (9,12,13). eIF4E phosphorylation was also well demonstrated to influence the translation of specialised mRNAs including those of IκBα that mediates type I IFN production (14).

mTOR is a S/T kinase that assembles into one of the two distinct complexes (mTORC1/mTORC2) (15,16). mTORC1 is a key regulatory hub for various cellular and physiological activities such as translation and autophagy (17,18). mTORC1 regulates protein translation through at least two major substrates, eIF4E binding protein (4EBP) and the ribosome S6 kinase, p70 S6K (19,20). mTORC1-mediated multi-site phosphorylation of 4EBP1 releases eIF4E from sequestration facilitating its availability for binding to 5’ caps of mature mRNAs. In spite of its perceivable key role in translation, mTORC1 inhibition does not result in total inhibition of translation activities (21–23). Studies in the past have identified several mRNAs that respond to mTORC1 activity. mRNAs with a 5’ terminal oligopyrimidine (5’ TOP) element are particularly vulnerable to translation inhibition caused by mTORC1 inhibitors such as rapamycin, Torin1 and pp242 (21,22,24,25).

eIF2, a heterotrimeric complex formed by the association of α, β, and γ subunits, is a critical component of the ternary complex with Met-tRNA^Met^ and GTP (eIF2-TC). eIF2-TC binds to the 40S ribosome to form 43S PIC (26,27). Of these, eIF2α is the regulatory subunit that upon phosphorylation at S52 leads to the inhibition of GDP exchange with GTP after the hydrolysis of the bound GTP. eIF2α is phosphorylated by integrated stress response (ISR) kinases HRI, GCN2, PKR, and PERK (26–28). Of the known pathways, eIF2α phosphorylation has the most significant impact on translation as this leads to the most severe translation inhibition (28).

Despite having their distinct spheres of regulation, there has not been any major understanding on how these pathways influence each other under specific stimuli barring a few reports (29). Certain degree of cross-activation of mTORC1 by ERK through p90 ribosomal S6 kinase- mediated phosphorylation of RAPTOR is reported (30). Partial dephosphorylation of eIF4E upon mTORC1 inhibition is also reported (31). Studies also cite certain amount of cross-talk between mTORC1 and eIF2α during autophagy (32). Interestingly, none of these pathways appear to wield a dominant role over the other. A recent study in fact reported that about 90% of the proteins affected by ISR induced by distinct signals are common barring a small subset of unique proteins (29). This also suggests the existence of alternate pathways to ensure basal translation activities under most of the conditions. Here, we report that co-inhibition of p38 MAPK and ERK1/2 resulted in dose-dependent phosphorylation of eIF2α indicative of ISR, strong dissociation of polysomes, and inhibition of translation. Both cap-dependent and cap-independent pathways were suppressed under these conditions. eIF2α phosphorylation by sodium arsenite induced p38 MAPK, which was critical in maintaining basal translation activities. Mnk1/2 inhibition failed to bring about similar effects indicating the involvement of alternate pathways downstream of the MAPKs leading up to eIF2α phosphorylation. Interestingly, Mnk1/2 inhibition activated Akt that could possibly have contributed to the stabilization of translation activities under this condition. Our studies demonstrate a dominant role for p38 and ERK1/2 in the regulation of protein translation, through eIF2α phosphorylation.

## Experimental procedures

### Cell culture, inhibitors, and antibodies

All the cell lines were cultured in DMEM supplemented with 10% Fetal Bovine Serum, penicillin-streptomycin and NEAA. Sodium arsenite, U0126, p38 MAPK inhibitor VIII, Akt-1/2 inhibitor VIII, and ETP-45835 were procured from Merck Millipore and Torin1 was purchased from Tocris Bioscience. All the primary antibodies except β-Tubulin and GAPDH (Thermo Fisher Scientific) were purchased from Cell Signaling Technology. HRP-conjugated secondary antibodies were from Jackson ImmunoResearch.

### Inhibitor treatments and immunoblotting

Cells were grown to 70-80% confluency, harvested, and lysed in NP-40 lysis buffer (33). Inhibitions were performed in most cases for 1-hour unless otherwise mentioned. Actively growing cells were treated with either vehicle or inhibitor-containing media before lysis. All the inhibitors were diluted in fresh DMEM right before the experiments. Protein concentration was estimated by BCA method (G Biosciences). Equal quantities of protein lysates were separated by SDS-PAGE, transferred to an activated PVDF membrane, and immunoblotting was performed as mentioned elsewhere (33). β-Tubulin and GAPDH were used as loading controls.

### Polysome preparation and profiling

Reagents for polysome analysis were purchased from Sigma and MP chemicals. Polysome profiles of the inhibited cells were analyzed as described earlier (33). Cells were grown in 150 cm^2^ flasks till 70% confluency and inhibition studies were performed for 1-hour. Following the inhibition, the cells were harvested and washed twice with ice-cold 1×PBS containing 100 μg/mL cycloheximide. The lysates were prepared in polysome lysis buffer (20 mM Tris-Cl pH 8.0, 140 mM KCl, 1.5 mM MgCl_2_, 0.5 mM DTT, 1% Triton X-100, 1× protease inhibitor, 0.5 mg/mL heparin, 100 μg/mL cycloheximide, 100 units of RNasin/mL) and crude RNA concentrations were spectrophotometrically measured. 175 μg of crude RNA lysates were layered on 11 mL of 10-50% linear sucrose gradients (20 mM Tris-Cl pH 8.0, 140 mM KCl, 1.5 mM MgCl_2_, 0.5 mM DTT, 100 μg/mL cycloheximide, 10mM PMSF, and 10-50% sucrose) and the resulting gradients were centrifuged in SW-41 Ti rotor (Beckman Coulter) at 35,000 r.p.m. at 4°C for 3-hours. The samples were fractionated using Teledyne ISCO fraction collector system with a constant monitoring of absorbance at 254 nm to generate Polysome profiles.

### Luciferase assay

Cells were seeded in 6 well plate format and grown till 70% confluency and the luciferase reporter plasmids were transfected using Lipofectamine 3000 (Thermo Fisher Scientific). One hour prior to harvesting, the transfected cells were treated with the specific inhibitors. Cells were harvested 10-hours post-transfection and the luciferase reads were quantified using Dual-Luciferase reagents (Promega) as per manufacturer’s protocol on EnSpire multimode plate reader (Perkin Elmer).

### OPP Assay

Total protein synthesis was quantified with the help of OPP-incorporation assay based on Click-iT™ chemistry using the Click-iT™ Plus OPP-Alexa Fluor™ 488 protein synthesis assay kit (Life Technologies). The assay was carried out in accordance with the manufacturer’s recommendations. Briefly, Huh7.5 cells were seeded in 8-well chamber slides (Lab-Tek) up to ~80% confluency, and subsequently treated with either DMSO, p38 MAPK inhibitor VIII, U0126 or both for 1-hour. Post-treatment, the cells were treated with 20 μM OPP for 30 minutes at 37°C, followed by fixing with 3.7% formaldehyde and permeabilization with 0.5% Triton X-100 in PBS. Click-iT reaction was carried out with Alexa Fluor 488-picolyl azide for 30 minutes followed by nuclear staining using NuclearMask™ Blue stain. Stained cells were imaged using AxioImager Z2 microscope using Alexa Fluor-488 and DAPI filters for detection. Threshold level was set using OPP-untreated cells. Images were captured at 20× magnification. Image analysis was done using FIJI. Mean whole-cell fluorescence of ~90 individual cells per sample from each set was measured after background subtraction from individual fields. Data is represented as mean fluorescence intensity.

### Cell viability assays

Cell viability assays for treatments were carried out using MTT or trypan blue exclusion assay. For all assays, cells were seeded till ~80% confluency and treated with the pharmaceutical compound as mentioned. Vehicle- or inhibitor-treated cells were trypsinized and mixed in 1:1 ratio with 0.4% trypan blue and counted using Neubauer chamber to determine the cell count. For MTT assay, media containing MTT (final concentration of 0.5 mg/mL) was added to cells post treatment and incubated at 37°C for 3.5-hours. Formazan crystals formed were dissolved in 500 μL DMSO with mild agitation for 30 minutes. The assay readout was measured as absorbance at 440 nm, with a reference reading at 650 nm.

### Statistical Analysis

Data from three independent experiments were used for statistical analysis using two-tailed Student’s *t*-test for viability and dual-luciferase assays and represented graphically as Mean ± SEM. One-way ANOVA was used for OPP assay analysis and plotted as median with interquartile range. *, ** and *** indicate *p*-values < 0.05, 0.005 and 0.0005, respectively.

## RESULTS

### p38 MAPK and ERK1/2 dual inhibition inhibit eIF4E phosphorylation in a dose-dependent manner

p38 and ERK1/2, two major MAPKs regulate the phosphorylation of eIF4E through phosphorylating and activating Mnk1/2 (3,5,34,35). However, there is little mechanistic evidence on their synergistic regulation of eIF4E and subsequent impact on translation. We addressed this question by comparing eIF4E phosphorylation status under conditions of individual and simultaneous inhibitions of both MAPKs. First, we compared the degree of inhibition in Huh7.5 cells upon three combinations of p38 MAPK inhibitor VIII (p38i) and U0126 (ERK1/2i), viz. 12.5/25 μM, 25/50 μM and 50/100 μM concentrations. The status of eIF4E phosphorylation after 1-hour inhibition was analyzed as the measure of inhibition. A gradual and dose-dependent dephosphorylation of Mnk1 and eIF4E was observed and total dephosphorylation was achieved at the combination of 50/100 μM concentrations (Figure 1A). However, cell viability was only modestly impacted by the treatments (Figure 1B). Total dephosphorylation of eIF4E was also achieved in MCF7 cells (Figure 1C). Next, we studied if the dual MAPK inhibition (dual-Mi) had any synergistic effect on eIF4E phosphorylation. Torin1, a potent mTOR inhibitor has been shown previously to affect translation. Torin1 treatment (750 nM) for 1-hour was done alongside MAPK inhibitions, to compare the effects of each treatment on MAPK substrate phosphorylation. Inhibition of mTORC1 was confirmed by the dephosphorylation of its substrates 4EBP1 and ULK1 (Figure 4A). As demonstrated in Figure 1D, the dual inhibition in Huh7.5 cells resulted in much stronger inhibition of Mnk1 and eIF4E than the individual inhibitions did, suggesting that p38 and ERK1/2 synergistically regulate eIF4E phosphorylation. No significant change was observed in the sample inhibited with Torin1, confirming the specificity of the treatments. Similar results were observed in MCF7 and HeLa cells (Figure S1 A & B, respectively)

**Figure 1.**
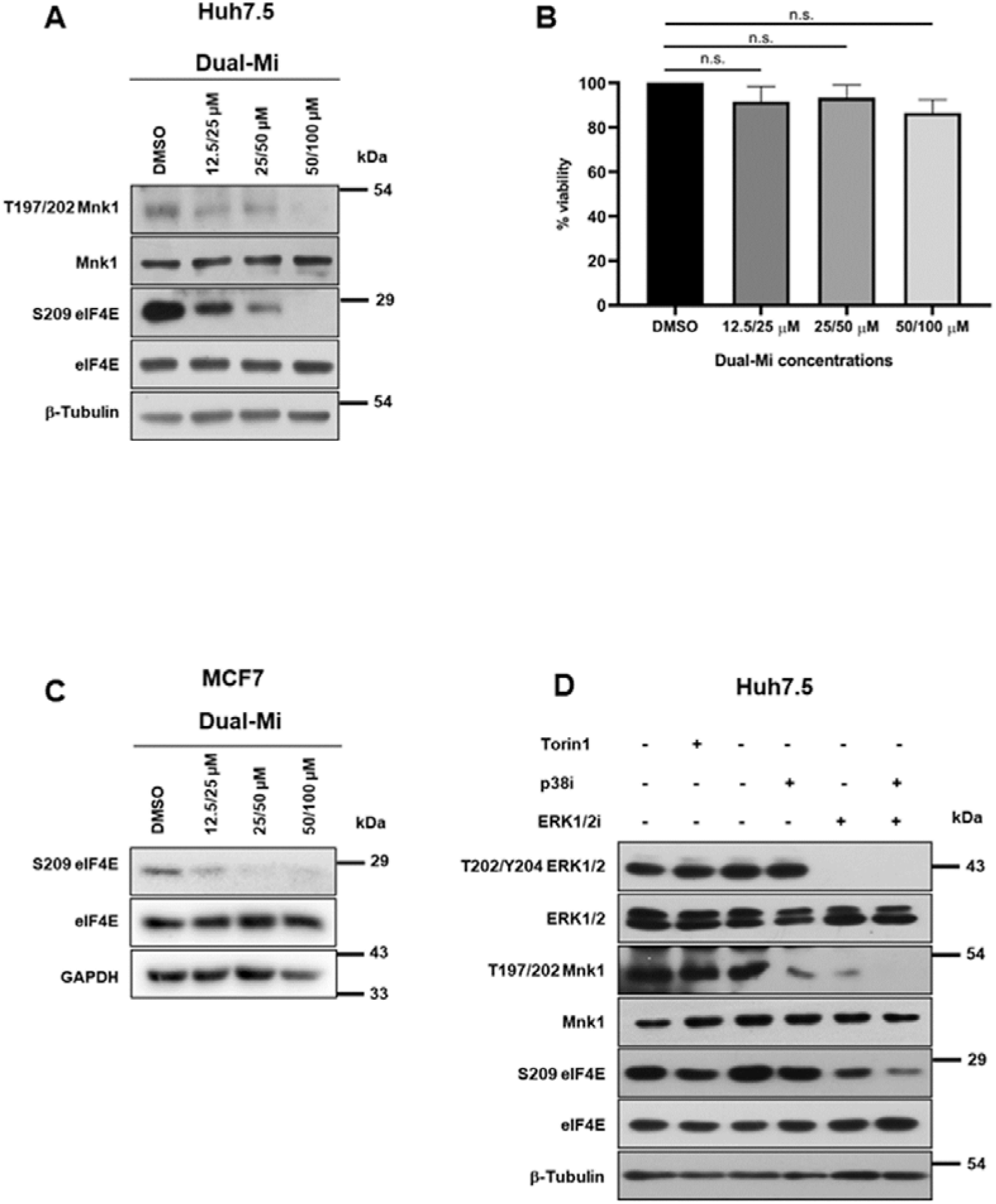
p38 and ERK1/2 MAPK co-inhibition causes dose dependent decrease in eIF4E phosphorylation. **(A)** Immunoblots depicting phosphorylation and expression status of Mnk1 and eIF4E from Huh7.5 cells upon treatment with 12.5/25, 25/50, or 50/100 μM of dual-MAPK inhibitors (Dual-Mi) for 1-hour along with DMSO vehicle control. (**B)** Huh7.5 cell viability was measured upon treatment with DMSO vehicle or different concentrations of inhibitors for 1-hour using trypan blue exclusion method. Graph is representative of 3 independent experiments. The statistical significance was calculated using two tailed, paired Student’s *t* test where n.s. represents non-significant. **(C)** Immunoblots depicting phosphorylation and expression of eIF4E from MCF7 cells upon similar treatment as in (A). **(D)** Immunoblots depicting phosphorylation and expression status of ERK1/2, Mnk1, and eIF4E from Huh7.5 cells treated with DMSO vehicle control, Torin1 (750 nM), p38i (50 μM), ERK1/2i (100 μM), or Dual-Mi (50/100 μM p38i/ERK1/2i). Lanes 1 and 3 are independent vehicle controls for Torin1 and MAPK inhibitors, respectively. p38i-p38 MAPK inhibitor VIII, ERK1/2i-U0126, Dual-Mi-p38i+ERK1/2i.

### Dual-Mi at high concentrations causes near-absolute collapse of polysomes and severe suppression of global translation

In order to characterize the effect of dual-Mi on global translation, polysome profiles in Huh7.5 cells treated with individual inhibitors or both (at 50/100 μM for p38 MAPK inhibitor VIII/U0126 respectively) were mapped. It may be noted that these concentrations were consciously chosen in this study so that the translation activities could be studied under conditions of complete inhibition of the MAPKs. Surprisingly, 1-hour dual-Mi in Huh7.5 cells caused near-absolute collapse of polysome peaks with corresponding accumulation of 80S, suggesting a global inhibition of translation activities (Figure 2A). No polysome peaks were visible under this condition, reminiscent of translation arrest caused by puromycin treatment (36). In comparison to dual-Mi, individual inhibitions brought about only modest effects on the polysome peaks (Figure 2B). The magnitude of polysome dissociation by dual-Mi was remarkably higher than those caused by eIF2α phosphorylation by various methods ((26,28) and Figure 6B) and mTORC1 inhibition by Torin1 (Figure 2C). Severe impact of dual-Mi on polysome association was observed in MCF7 cells as well (Figure S2A). Moderate collapse of polysomes was visible at a lower concentration of 25/50 μM (Figure S2B), once again underlining the specificity of the response to the inhibitions. The results demonstrated that the concurrent loss of activity of the two MAPKs severely inhibits polysome assembly and possibly translation.

**Figure 2.**
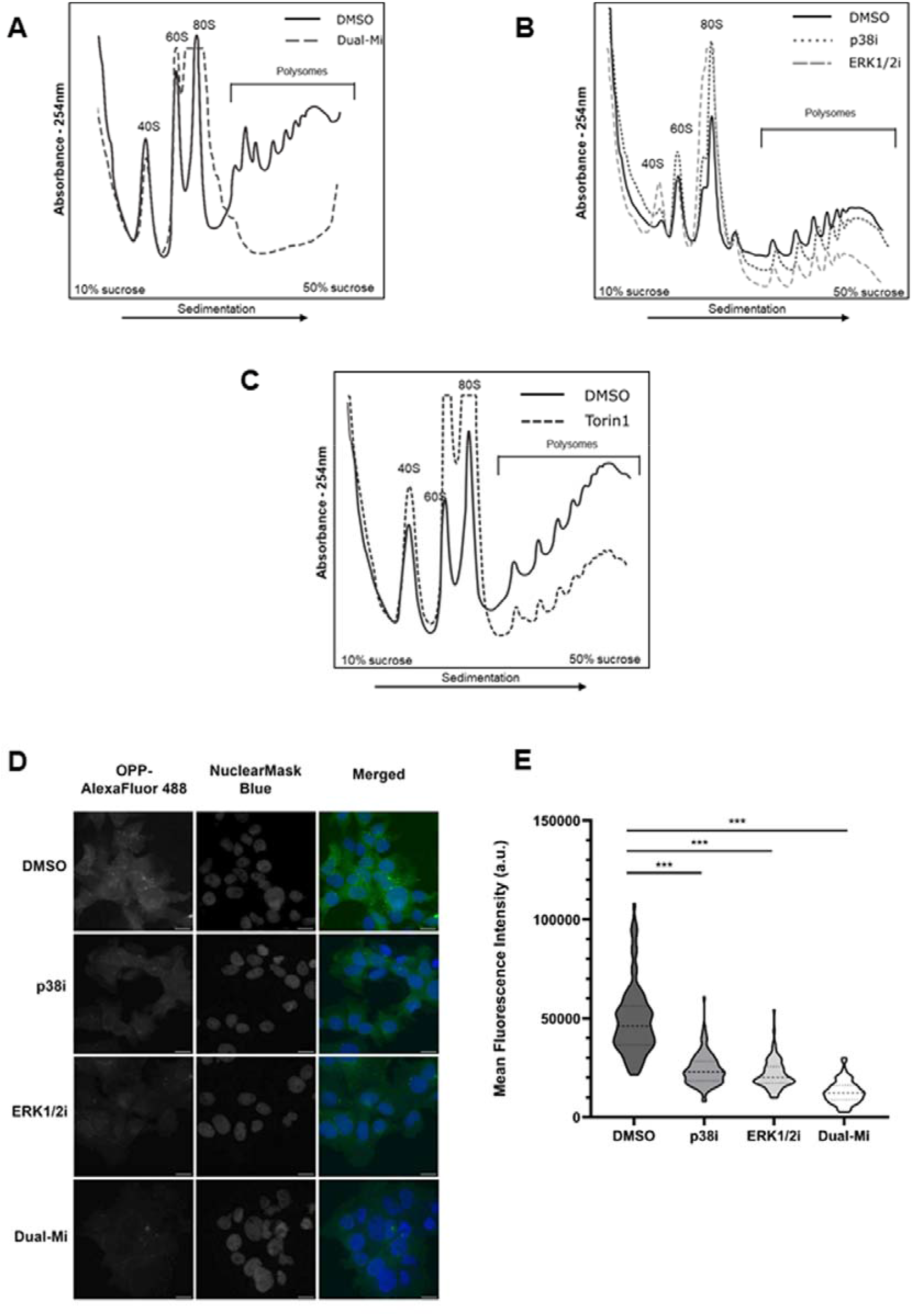
p38 and ERK1/2 MAPK dual inhibition severely affects polysome stability and suppresses translation. **(A-C)** Polysome profiles of Huh7.5 cells treated with DMSO vehicle control or specific inhibitor(s) for 1-hour. Polysome profile analyses were performed after density gradient ultracentrifugation of the corresponding cytosolic extract. Free ribosomal subunits (40S and 60S), monosome (80S) and the polyribosomes were fractionated by measuring the absorbance at 254 nm. Each graph shows treatment-curve overlaid on the vehicle control. **(A)** From dual-Mi (50/100 μM p38i/ERK1/2i) treatment. **(B)** From p38i (50 μM) and ERK1/2i (100 μM) individual treatments. **(C)** From Torin1 (750 nM) treatment. **(D)** Representative images from OPP-incorporation assay for assessing protein synthesis in DMSO, p38i (50 μM), ERK1/2i (100 μM) and dual-Mi (50/100 μM p38i/ERK1/2i), performed in Huh7.5 cells. AlexaFluor 488 conjugated-OPP (green) was used to determine level of nascent protein synthesis. NuclearMask Blue (blue) was used to stain the nucleus. Images were captured at 20× magnification and the scale bar represents 20 μm length. **(E)** Quantitative analysis of OPP-incorporation depicted as violin plots. Data represented is from ~90 cells as 3 independent experimental setups and is represented with median values and minimum and maximum quartiles, and *p*-value was calculated using one-way ANOVA; *** represents *p* < 0.0005.

We next quantified the translation activities during dual-Mi in Huh7.5 cells by labelling the nascent polypeptides using O-propargyl-puromycin (OPP) incorporation assay. Cells were treated with p38 or ERK1/2 inhibitor or both of them for 1-hour and were labeled with OPP for 30 minutes. Subsequently, they were analyzed by fluorescence microscopy to quantify protein synthesis. As demonstrated in Figure 2D, appreciable fluorescence was detected throughout cytoplasm accompanied by several bright foci in control cells indicating active translation. Both p38 and ERK1/2 inhibitions decreased the incorporation of OPP, evident from both lower intensity fluorescence and reduced number of bright foci, implying lower translation rates. The strongest inhibition of OPP incorporation was detected in dual-Mi treated cells indicating very low levels of translation in them. Quantitative analysis of fluorescence intensities revealed approximate intensity drop of 50% in samples treated with either of the inhibitors and about 75% drop in dual-Mi samples, confirming strong inhibition of translation activities in these cells (Figure 2E), substantiating the polysome dissociation. These results also demonstrate that p38 and ERK1/2 are crucial guardians of eukaryotic translation.

### Mnk1/2 inhibition and eIF4E dephosphorylation do not recapitulate the polysome collapse caused by dual-Mi

Though Mnk1/2 double KO mice did not display any translation and growth defect, studies have demonstrated Mnk1/2 as key molecules in the phosphorylation of eIF4E (12). We studied if the effects on translation during dual-Mi are channelled through Mnk1/2. Mnk1/2 inhibitor ETP-45835 brought about eIF4E dephosphorylation in a dose-dependent manner (Figure 3A). In agreement with previous reports, polysome profiles of Mnk1/2 inhibited cells did not demonstrate any appreciable level of polysome dissociation (Figure 3B) in contrast to dual-Mi treatment (Figure 2A). These results indicate the participation of other signal pathways regulating polysome assembly and translation during dual-Mi treatment.

**Figure 3.**
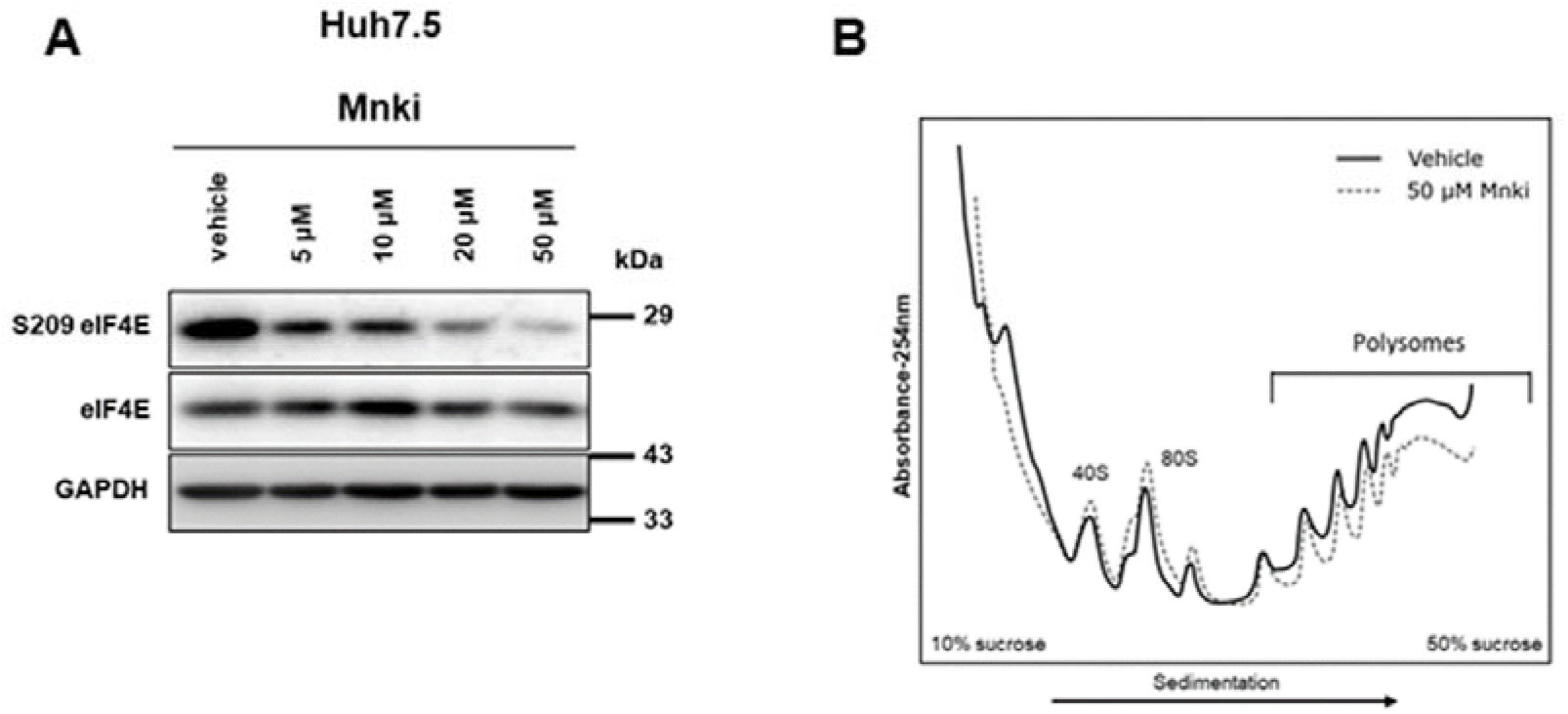
Polysome collapse caused by dual-Mi is independent of Mnk1/2 inhibition and eIF4E dephosphorylation. **(A)** Immunoblots depicting phosphorylation and expression status of eIF4E from Huh7.5 cells upon treatment with 5, 10, 20 or 50 μM of Mnki for 1-hour along with water vehicle control. (**B)** Polysome profiles from Huh7.5 cells upon treatment with vehicle or Mnki (50 μM) for 1-hour. Free ribosomal subunits (40S and 60S), monosome (80S) and the polysomes were fractionated by measuring the absorbance at 254 nm after density gradient centrifugation of corresponding cytosolic extracts. The graph shows treatment-curve overlaid on the vehicle. Mnki - ETP-45835

**Figure 4.**
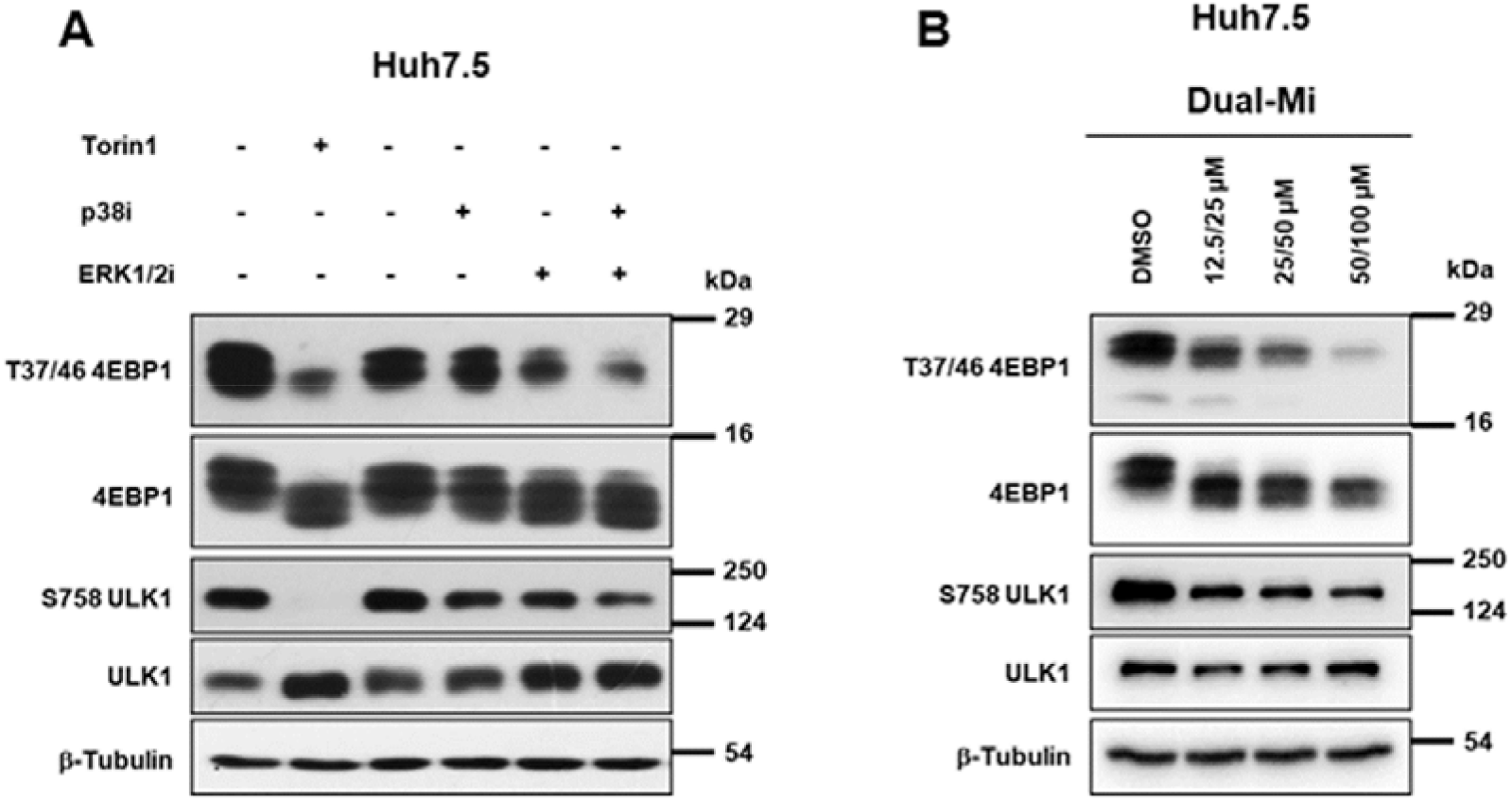
Dual-Mi causes moderate inhibition of mTORC1 activity in Huh7.5. **(A)** Immunoblots depicting phosphorylation and expression status of 4EBP1 and ULK1 from Huh7.5 cells treated with DMSO vehicle control, Torin1 (750 nM), p38i (50 μM), ERK1/2i (100 μM), or Dual-Mi (50/100 μM p38i/ERK1/2i) for 1-hour. (**B)** Immunoblot depicting phosphorylation and expression status of 4EBP1 and ULK1 from Huh7.5 cells upon treatment with 12.5/25, 25/50, or 50/100 μM of p38i/ERK1/2i dual-MAPK inhibitors for 1-hour along with DMSO vehicle control.

### Dual-Mi causes moderate inhibition of mTORC1 activity

Since eIF4E dephosphorylation caused no significant impact on the polysome assembly, we studied the involvement of the other known pathways in coordinating the polysome dissociation by dual-Mi. mTORC1 pathway is a well-known regulator of translation and its inhibition causes significant depletion of polysomes and loss in translation activities ((22) and Figure 2C). We analyzed whether dual-Mi causes mTORC1 inhibition. Huh7.5 cells displayed moderate inhibition of mTORC1 activity, evident from the dephosphorylation of 4EBP1 and ULK1, two important substrates of mTORC1 (Figure 4A). Dose-dependence of the inhibition demonstrated the specificity of this observation (Figure 4B). Individually, ERK1/2 inhibition induced a stronger mTORC1 inhibition than p38 inhibition (Figure 4A). Torin1, an mTOR inhibitor, caused severe inhibition of mTORC1 activity. The degree of mTORC1 inactivation was less remarkable in MCF7 and HeLa cells (Figure S3 A & B, respectively), indicating that this effect on mTORC1 activity is cell-specific and hence may not be contributing to the polysome dissociation by dual-Mi.

### Dual-Mi induces strong eIF2α phosphorylation in a dose-dependent manner

Next, we tested the effect of dual-Mi on eIF2α phosphorylation. Huh7.5 cells subjected to dual-Mi were analyzed for the status of eIF2α phosphorylation. Individually, p38 inhibition caused a moderate eIF2α phosphorylation while ERK1/2 inhibition caused a stronger phosphorylation (Figure 5A). Interestingly, dual-Mi brought about a much stronger phosphorylation than the individual inhibitions, indicating that the two MAPKs regulate eIF2α independent of each other. These results also suggested that eIF2α phosphorylation through ISR could be a critical event behind the polysome dissociation during dual-Mi. A dose-dependent phosphorylation of eIF2α in response to varying concentrations of dual-Mi confirmed the specificity of the response (Figure 5B). eIF2α phosphorylation was consistently observed in MCF7 and HeLa cells as well (Figure S4 A & B respectively). Our results demonstrate that p38 and ERK1/2 are important players in the regulation of eIF2α phosphorylation and in the translation activities. Mnk1/2 inhibition could not recapitulate eIF2α phosphorylation (Figure 5C), ruling out its participation in the translation suppression associated with dual-Mi.

**Figure 5.**
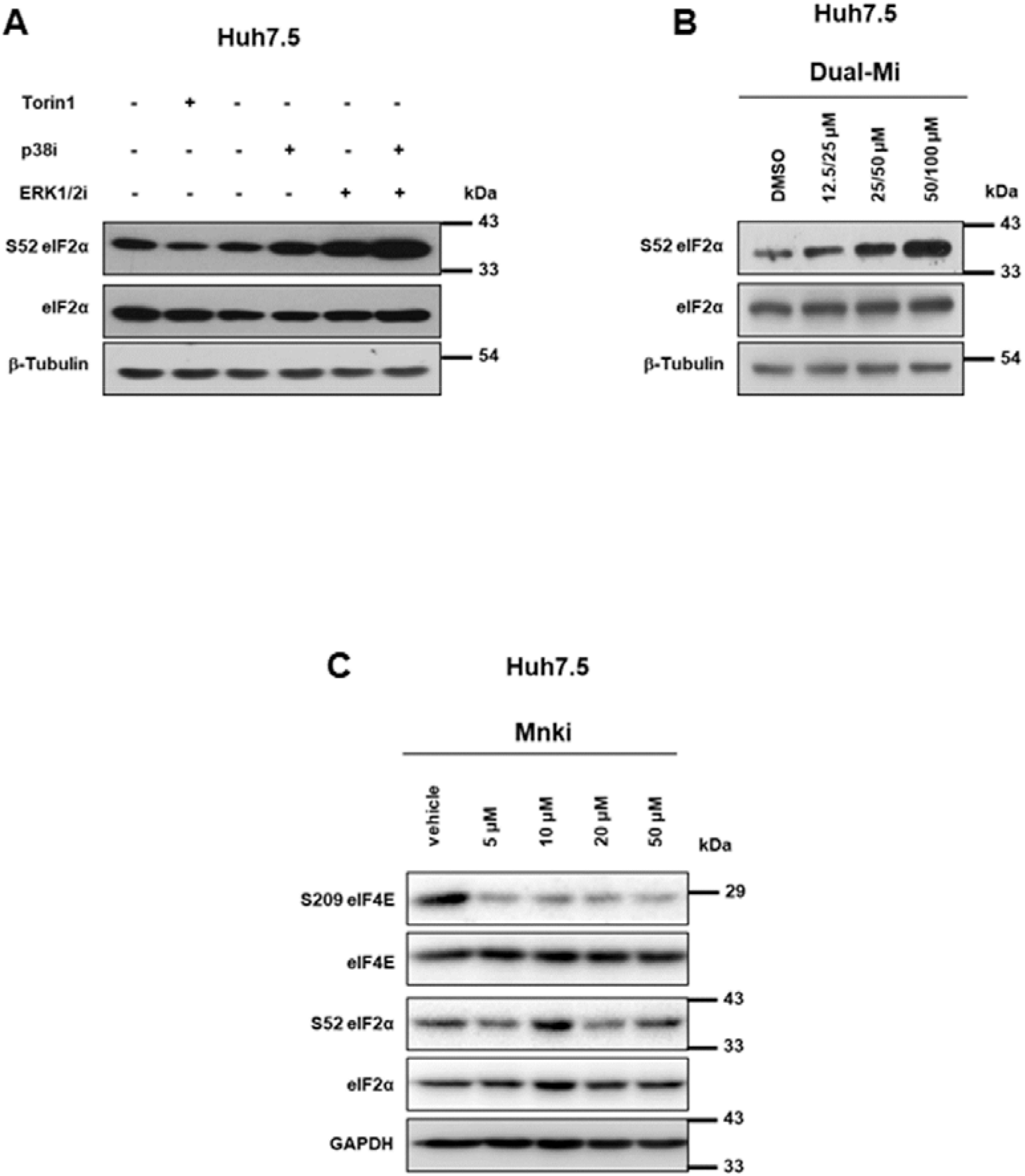
p38 and ERK1/2 MAPKs synergistically regulate eIF2 α phosphorylation independent of its common downstream kinase Mnk1/2. **(A)** Immunoblots depicting phosphorylation status of eIF2α from Huh7.5 cells treated with DMSO vehicle control, Torin1 (750 nM), p38i (50 μM), ERK1/2i (100 μM) or Dual-Mi (50/100 μM p38i /ERK1/2i) for 1-hour. **(B)** Immunoblots depicting phosphorylation and expression status of eIF2α from Huh7.5 cells upon treatment with 12.5/25, 25/50, or 50/100 μM of Dual-Mi for 1-hour along with DMSO vehicle control. **(C)** Immunoblots depicting phosphorylation of eIF4E and eIF2α from Huh7.5 cells upon treatment with 5, 10, 20 or 50 μM of Mnki for 1-hour along with water vehicle control.

### Activation of p38 upon stress is pivotal for the maintenance of low-level translation during eIF2α phosphorylation

The role of p38 as a key molecule facilitating stress response is well documented (37). We investigated the role of the two MAPKs in response to stress induced by general translation inhibitor sodium arsenite. Arsenite activates ISR kinases HRI, PKR, PERK and GCN that phosphorylate eIF2α (38–41). Treatment of Huh7.5 cells with 40 μM arsenite induced strong eIF2α phosphorylation as expected. Interestingly, p38, but not ERK1/2 was phosphorylated by the treatments (Figure 6A), indicating its crucial role in the cell survival upon translation attenuation during ISR. Similar results were observed in MCF7cells (Figure 6A). Intriguingly, stress-induced p38 phosphorylation did not cause Mnk1/2 and eIF4E phosphorylation (Figure 6A), indicating the involvement of other mechanisms in the regulation of eIF4E phosphorylation by p38 under conditions that promote it. Sodium arsenite did not affect mTORC1 substrates in both Huh7.5 and MCF7 cells, ruling out the possibility of mTORC1 participating in ISR (Figure S5 A & B, respectively). Polysome analysis of arsenite treated cells showed a drop in translation as expected (Figure 6B). Despite a strong collapse of the heavy polysomes, lighter polysomes were intact. As indicated earlier, this polysome dissociation was less remarkable than that by dual-Mi, indicating the stronger translation inhibitory effects by the latter (Figures 2A & 6B).

**Figure 6.**
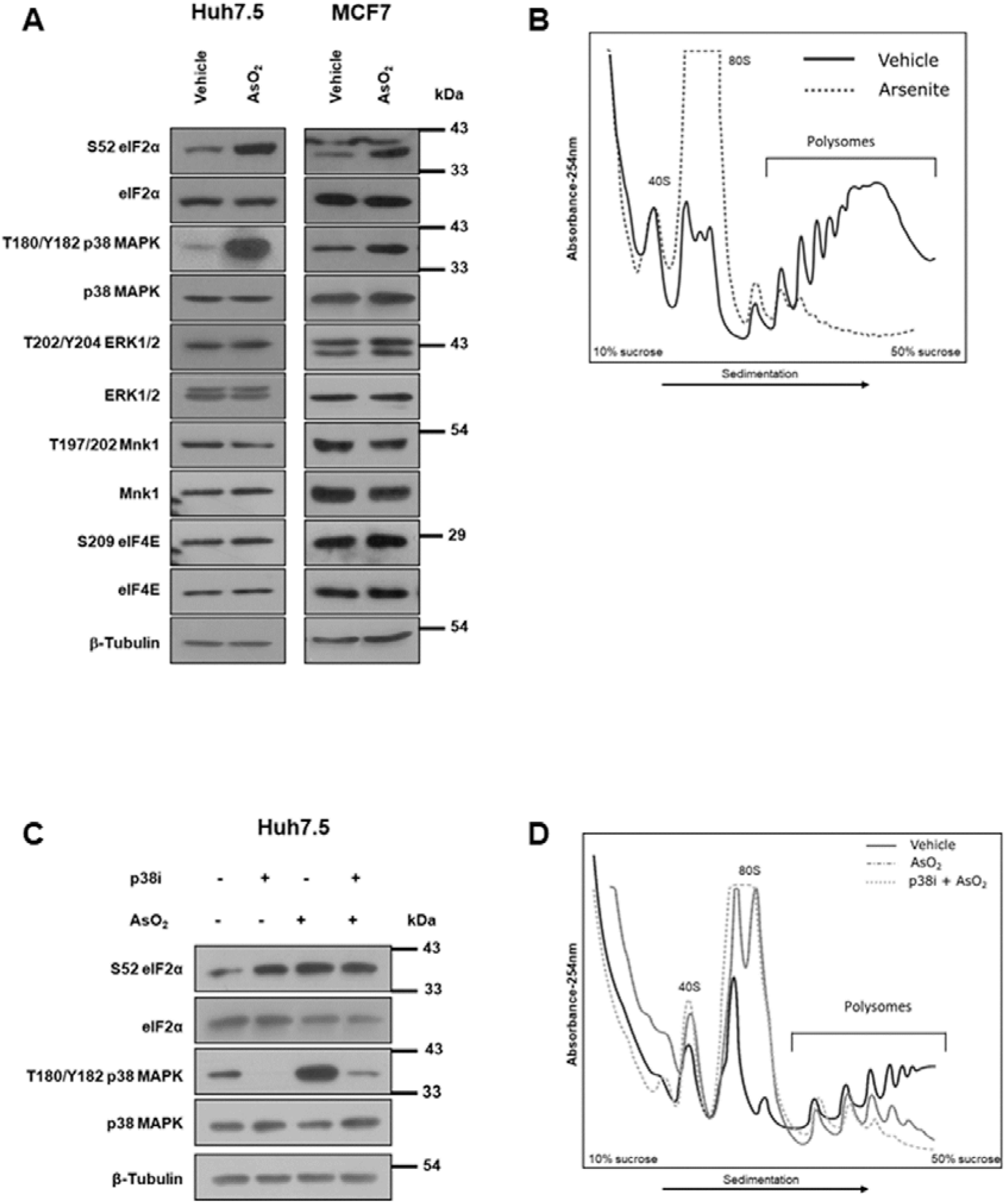
Arsenite induced p38 MAPK activation is critical for polysome stability. **(A)** Immunoblots depicting phosphorylation and expression status of eIF2α, p38 MAPK, ERK1/2, Mnk1 and eIF4E from Huh7.5 and MCF7 cells upon treatment with vehicle or sodium arsenite (40 μM) for 1-hour. **(B)** Polysome profiles from Huh7.5 cells treated with vehicle or sodium arsenite (40 μM) for 1-hour. Free ribosomal subunits (40S and 60S), monosome (80S) and the polyribosomes were fractionated by measuring the absorbance at 254 nm after density gradient centrifugation of corresponding cytosolic extracts. The graph shows treatment-curve overlaid on that of the vehicle. **(C)** Immunoblots depicting phosphorylation and expression status of eIF2α and p38 MAPK upon treatment with vehicle, sodium arsenite (40 μM), p38i (50 μM) or both (40/50 μM sodium arsenite/p38i) for 1-hour. **(D)** Polysome profiles from Huh7.5 cells treated with vehicle, sodium arsenite (40 μM), or both (40/50 μM sodium arsenite/p38i) for 1-hour. Free ribosomal subunits (40S and 60S), monosome (80S) and the ribosomes were fractionated by measuring the absorbance at 254 nm after density gradient centrifugation of corresponding cytosolic extracts. The graph shows treatment-curve overlaid on the vehicle. AsO_2_-Sodium arsenite.

Translation arrest is a general response to ISR and our results suggest that p38 activation is a probable feedback mechanism to sustain translation. In order to characterize the role of activated p38 upon ISR, we treated Huh7.5 cells with arsenite along with p38 inhibitor for 1-hour. This would, in principle, disallow the activation of p38 during ISR and hence could be an ideal set up to study the role of p38 in translation during ISR. Independently, p38 inhibition induced eIF2α phosphorylation similar in magnitude as arsenite did and their combined treatment did not further induce it (Figure 6C). Quite interestingly and as hypothesized, inhibition of p38 in cells treated with arsenite caused more severe polysome collapse as compared with arsenite treatment alone (Figure 6D). This result indicates that p38 activation is a crucial and remedial consequence to ISR in the maintenance of minimal translation activity required for the synthesis of necessary proteins. Our results demonstrate that activation of p38 is crucial in sustenance of translation upon ISR induction.

### Both cap-dependent and independent translations are inhibited by dual-Mi

Even though the default mode of translation is 5’-cap-dependent, cap-independent translation is promoted under several physiological conditions of stress (42). Even under eIF2α phosphorylation mediated translational arrest, a set of mRNAs are translated actively. To verify whether dual-Mi suppresses cap-independent translation, we used a bicistronic luciferase assay where translation of *Renilla* luciferase (Rluc) is cap-dependent whereas that of firefly luciferase (Fluc) is driven by HCV or EMCV IRES (Figure 7A) (43). Cells transfected with either of these vectors were subjected to dual-Mi and translation efficiencies were measured through luciferase activities. Interestingly, both Fluc and Rluc activities from HCV IRES construct were inhibited by approximately 60% after 1-hour of inhibition, indicating that dual-Mi mediated translation arrest inhibits both cap-dependent and cap-independent translation (Figure 7B). Inhibitions of similar magnitude from EMCV IRES construct confirmed the above observation (Figure 7C). A conventional analysis based on F/R ratio would not identify these inhibitions as both cap-dependent and independent translations were impacted similarly. These results are in agreement with the severe translation arrest described in previous section and indicate that concurrent inhibition of p38 and ERK1/2 affects translation machinery as a whole.

**Figure 7.**
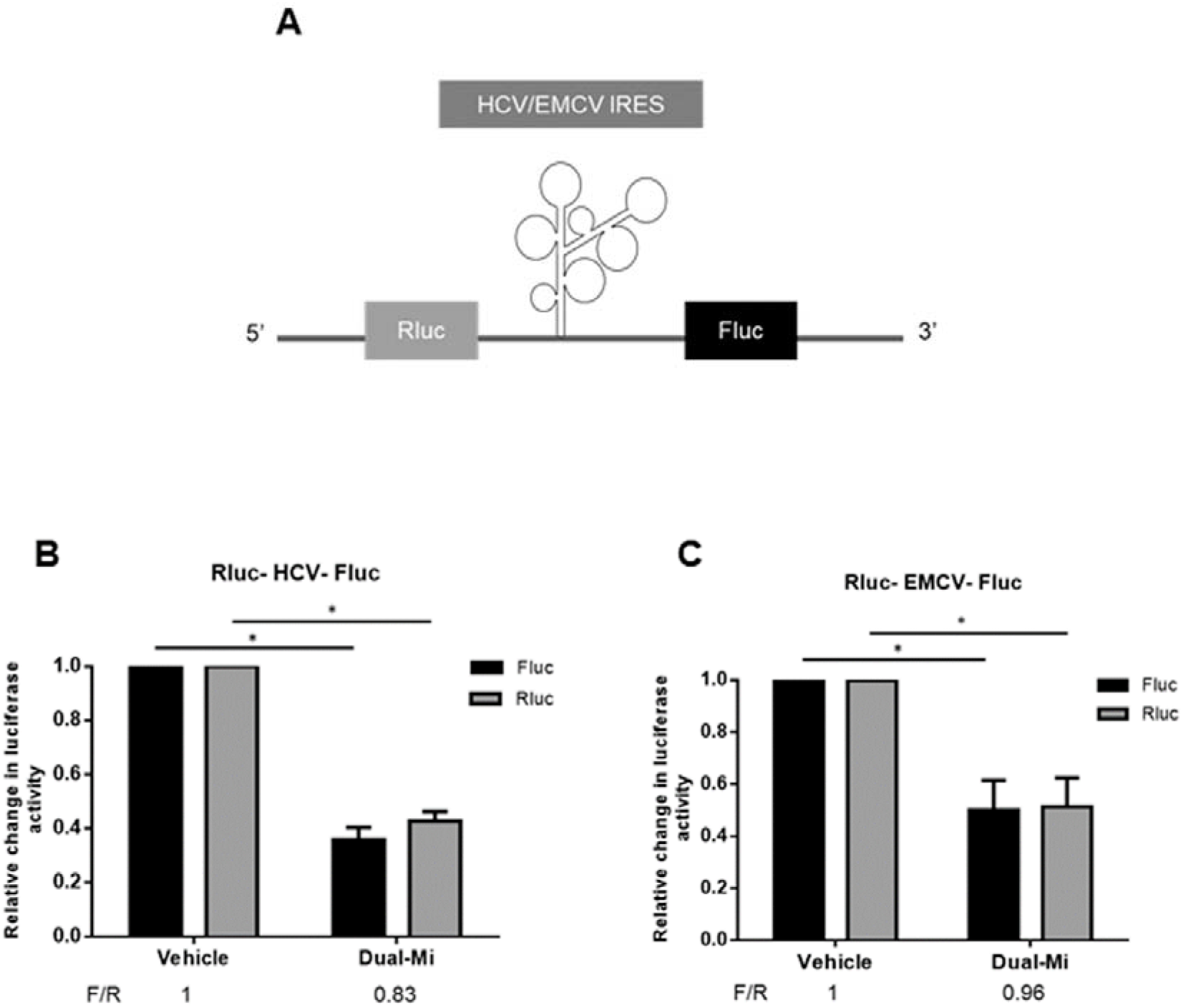
Dual-Mi inhibits both cap-dependent and cap-independent translation. **(A)** Schematic representation of dual-luciferase reporter construct where HCV/EMCV IRES elements are sandwiched between *Renilla* (Rluc) and Firefly (Fluc) luciferase genes. (**B and C)** Luciferase assay in Huh7.5 lysates transfected with HCV **(B**) and EMCV **(C)** dual-luciferase reporter plasmids for 9-hours and further treatment either with vehicle or dual-Mi (50/100 μM p38i/ERK1/2i) for 1-hour. Relative change in luciferase activity was calculated by normalizing Fluc/Rluc reading of inhibitor treated cells with its corresponding vehicle control. *p*-value was calculated using two tailed, paired Student’s *t*-test; * represents *p* < 0.05. F/R ratio of each treatment is represented below each graph.

### eIF2α phosphorylation is reversed during long-term dual-Mi

The profound loss in polysome assembly and translation inhibition during dual-Mi for 1-hour did not appear to affect the cellular viability. Long-term dual-Mi significantly inhibited cell viability as anticipated (Figure 8A), but, a considerable population of cells remained active at the end of the treatment. We investigated if the ISR is reversed during the long-term dual-Mi (24-hours) in Huh7.5 cells. eIF4E remained inhibited throughout the inhibition period, confirming the sustained inhibition of the MAPKs (Figure 8B). Interestingly, eIF2α remained phosphorylated until 12-hours post treatment and subsequently returned to the basal levels (Figure 8B), indicating that the ISR was reversed by 24-hours. Since Akt is an important regulator in cell survival (44), we studied its phosphorylation. The surviving cells indeed displayed higher phosphorylation of Akt from 4-hours of treatment onwards and gradually increased until 24-hours (Figure 8C). These results demonstrate that despite a severe inhibition of translation activities by dual-Mi, a significant fraction of the cells recover from this inhibition over a period time and we speculate that Akt could be an important player in this survival.

**Figure 8.**
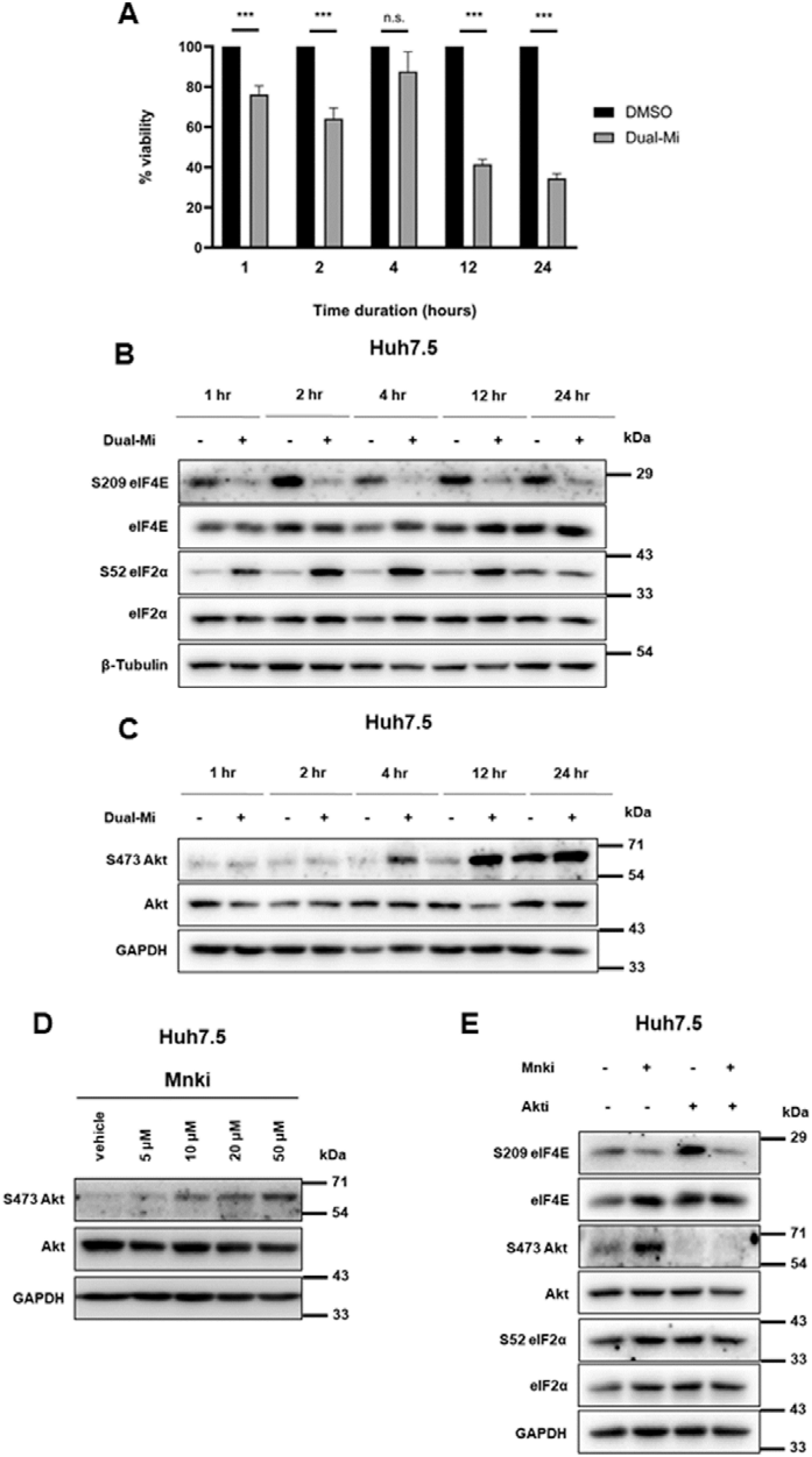
Chronic inhibition of p38 and ERK1/2 MAPK results in activation of alternate cell survival pathways to support cell sustenance. **(A)** Viability of Huh7.5 cells upon treatment with DMSO vehicle control or dual-Mi (50/100 μM p38i/ERK1/2i) for 1-, 2-, 4-, 12- or 24-hours using MTT assay. Graph is representative of three independent experiments. *p*-value was calculated using two tailed, paired Student’s *t*-test; where *** represents *p* < 0.0005 and n.s. represents non-significant. **(B)** Immunoblots depicting phosphorylation and expression status of eIF2α, eIF4E from Huh7.5 cells upon treatment with DMSO vehicle control or dual-Mi (50/100 μM p38i/ERK1/2i) for 1-, 2-, 4-, 12- or 24-hours. **(C)** Those of Akt from the same treatment as in (B). **(D)** Immunoblots depicting phosphorylation and expression status of Akt from Huh7.5 cells upon treatment with 5, 10, 20 or 50 μM of Mnki for 1 hour along with water vehicle control. **(E)** Immunoblots depicting phosphorylation and expression status of eIF4E, Akt and eIF2α from Huh7.5 cells upon treatment with vehicle, Mnki (10 μM), Akti (5 μM) or both (10/5 μM Mnki/Akti), Akti-Akt inhibitor VIII.

### Mnk1/2 inhibition promotes Akt phosphorylation

While investigating the potential mechanism of activation of Akt phosphorylation during the prolonged dual-Mi, we noticed that Mnk1/2 inhibition for 1-hour resulted in higher S473 Akt phosphorylation in a dose-dependent manner (Figure 8D) indicating that Mnk1/2 inhibition is inducing mTORC2. As in the case of Mnk1/2 (Figure 5C), the concurrent inhibitions of Akt and Mnk1/2 failed to make any change in the phosphorylation status of eIF2α (Figure 8E), ruling out the involvement of Akt during dual-Mi. However, it is compelling to suggest that Akt phosphorylation during prolonged dual-Mi could be important in the survival of the cells.

## DISCUSSION

Being a complex process, translation is often studied from single-pathway perspectives. Recent advancements in RNA deep sequencing and ribosome profiling has revolutionized the field by enabling identification of the transcripts that are regulated by distinct signal pathways at multiple stages of the process. However, the cross talks between major pathways involved in the translation regulation need deeper understanding. A recent comprehensive report explained cross-talk between mTORC1 and eIF2α pathways (29). Another report suggests that upon DNA damage, mTORC1 may regulate eIF2α phosphorylation via PERK and GCN2, to promote cell migration (45). Yet another study proposes that HRI-mediated ISR may inhibit mTORC1 activity in the liver to mitigate ineffective erythropoiesis (46). In these contexts, our study reveals a central role of p38 and ERK1/2, two MAPKs that have been shown to influence translation initiation by phosphorylating eIF4E. Our study identifies a novel mechanism of regulation where p38 and ERK1/2 regulate eIF2α phosphorylation and hence ISR. The impact of dual inhibition was profound and achieved near-total polysome dissociation that is not detected in any other individual conditions including mTOR inhibition and eIF2α phosphorylation by known agents. Clearly, the two molecules synergistically regulated eIF2α. Even as this study has not identified the specific kinase that mediates eIF2α phosphorylation, it helps in characterizing a major mechanism that has not been reported before.

Our studies underline the importance of p38 in maintaining the translation homeostasis upon stress (47). p38, but not ERK1/2, was activated upon arsenite treatment, indicating that the former is important in conditions of ISR. Interestingly, p38 phosphorylation did not lead to eIF4E phosphorylation suggesting the requirement of other factors in the regulation of the cap-binding protein. Additionally, ERK1/2 inhibition caused more noticeable dephosphorylation of eIF4E in all cell lines tested, implying the stronger influence by this MAPK. Although p38 activation upon arsenite treatment has been documented previously (48,49), our interests were to investigate its role in maintenance of translation under conditions of arsenite toxicity. Treatment of cells with arsenite and p38i simultaneously revealed a further depletion of polysomes, lending additional proof to its activation status under conditions of arsenite stress. Interestingly,

The most studied target of the two MAPKs in translation regulation is Mnk1/2. Previous studies have looked very closely at the role of Mnk1/2 and eIF4E phosphorylation in translation and their impact on global translation was less than appealing (50,51). Neither Mnk1/2 double knockout nor specific inhibitors have identified any significant impact on polysome assembly (12,52). Our studies using Mnk1/2 inhibitor are in total agreement with this. Importantly, Mnk1/2 inhibition did not bring about eIF2α phosphorylation unlike dual-Mi did, thereby excluding its participation in the mechanism we describe. It would be interesting to characterize the involvement of the other known downstream targets of MAPKs, RSK and MAPKAPKs under similar conditions.

Despite playing a major role, eIF2α phosphorylation alone cannot be implicated in the magnitude of translation arrest by dual-Mi. That is because eIF2α phosphorylation by various agents does not trigger such collapse in polysome assembly as in dual-Mi as demonstrated by our study and others (38,53,54). Thus dual-Mi appears to involve additional pathways and enforce a consolidated effect on translation. mTORC1 does not appear to be a key player in this as it was not inhibited consistently across cell types during dual-Mi.

We considered whether Akt phosphorylation upon Mnk1/2 inhibition protects eIF2α from phosphorylation. However dual inhibition of Mnk1/2 and Akt failed to notice any change. Interestingly, dual-Mi prevented Akt phosphorylation for long duration further strengthening the possibility of Akt phosphorylation in the process. However, appearance of phospho-Akt from 4-hours onwards indicated its possible role in the diminishing effects of dual-Mi and revival of the cells. As an upstream effector of the mTOR pathway that senses nutrient and oxygen deprivation and as a pro-survival signal, activation of Akt is an intriguing observation that suggests a possible feedback mechanism that initiates at later time points in dual-Mi.

Specificity and universal appeal of our findings were strengthened by the consistency of the observations across three distinct cell lines. The original objective of this study was to investigate the impact of total dephosphorylation of eIF4E on the polysome association and hence we consciously chose higher concentrations of MAPK inhibitors. eIF2α phosphorylation, the key finding in this study, was induced even at lower concentrations of inhibitors and followed a dose-dependency indicating the specificity of their effect. In addition, the cell viability was only moderate at 1-hour inhibition when most of our studies were performed.

Dual-Mi had prolonged effect on eIF2α phosphorylation. The inhibitors caused phosphorylation as long as 12-hours indicating that feedback mechanisms to neutralize ISR were not effective. One major question is how the cells are able to sustain without crashing during a severe translational repression. Quite clearly, a set of proteins critical for survival were being translated despite very low translation activities. It would be interesting to identify such proteins and the mechanisms that allow their translation. Activation of cap-independent mechanism is quite well accepted under conditions of translation suppression by mTORC1 inhibition (55). However, since dual-Mi caused severe suppression of cap-independent translation as well, the translating proteins are less likely to use this mode of translation. ERK1/2 activation promotes cell survival and proliferation in response to growth stimuli by driving the expression of pro-survival proteins. Response of p38 to stress stimuli depends more on the kinetics of its activation and can thus be either pro-survival or pro-apoptotic (56). Comparing our long duration dual-Mi observations to current understandings of MAPK signalling, it is possible that this cross-talk between MAPKs in general is allowing cells to remain viable despite seemingly negligible amounts of translation in the presence of high levels of eIF2α phosphorylation.

All MAPKs are known to respond to ER-stress in a myriad of ways, from transcriptionally upregulating pro-survival molecules, to stemming apoptotic signals, and seem to behave differently in different cell lines. p38 activation during ER-stress has been shown to cause switch from autophagy to apoptosis, mediated by PERK and eIF2α (57). MEK-ERK signalling has also been shown to be necessary for combating amino-acid deprivation in hepatocytes through GCN2 activation (58). These studies primarily see MAPK activation as a response to ISR that help combat the stress. Our studies also demonstrate that p38 is critical in the basal translation activities during stress. However, we also demonstrate that inhibition of these MAPKs can cause severe ISR. From our studies, p38 appears to be more critical to the balance in translation activities. Though inhibition of either of the two MAPKs caused eIF2α phosphorylation, only p38, not ERK1/2, was activated upon ISR induction. Since p38 inhibition causes ISR and eIF2α phosphorylation, we speculate that the subsequent feedback phosphorylation of p38 could be critical in limiting the ISR and stabilizing translation. Thus p38 seems to be a critical molecule in the post-ISR rescue of translation activities. Additionally, p38 could also be critical in maintaining the low translation activities during ISR.

## Author contributions

S.P., H.P., and K.H.H., conceived the study. S.P., H.P., D.V., D.G., and H.G.N performed the experiments. S.P., H.P., and K.H.H., analyzed the results. K.H.H wrote the manuscript while S.P., and H.P., edited it.

## Acknowledgement

We thank Rupesh Balaji for assisting in polysome profiling. Special thanks to Mohan Singh for logistical assistance for several experiments. HCV-IRES and EMCV-IRES constructs were kind gifts from Dr. Saumitra Das.

## Funding

This work was supported by funding from Department of Biotechnology, Govt. of India (BT/PR21356/MED/30/1779/2016). S.P, H.P and D.G received fellowships from Council of Scientific and Industrial Research, Govt. of India.

## Supplementary information

**Supplementary Figure S1.**
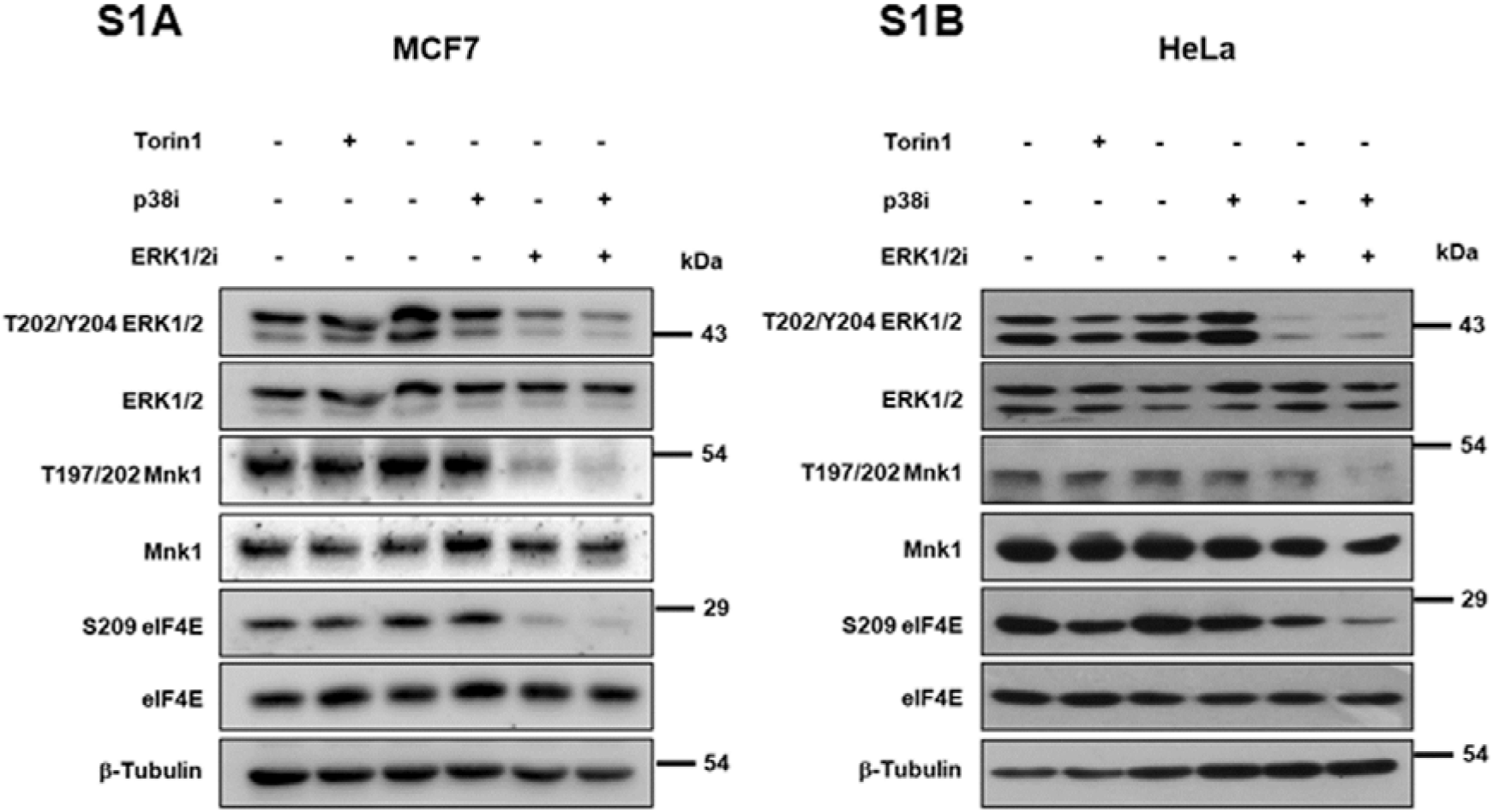
p38 and ERK1/2 MAPK synergistically regulates eIF4E phosphorylation across multiple cell lines. **(S1A)** Immunoblots depicting phosphorylation and expression status of ERK1/2, Mnk1 and eIF4E from MCF7 cells treated with DMSO vehicle control, Torin1 (750 nM), p38i (50 μM), ERK1/2i (100 μM) or Dual-Mi (50/100 μM p38i/ERK1/2i). **(S1B)** Immunoblots depicting phosphorylation and expression status of ERK1/2, Mnk1, and eIF4E from HeLa cells treated as mentioned in (S1A).

**Supplementary Figure S2.**
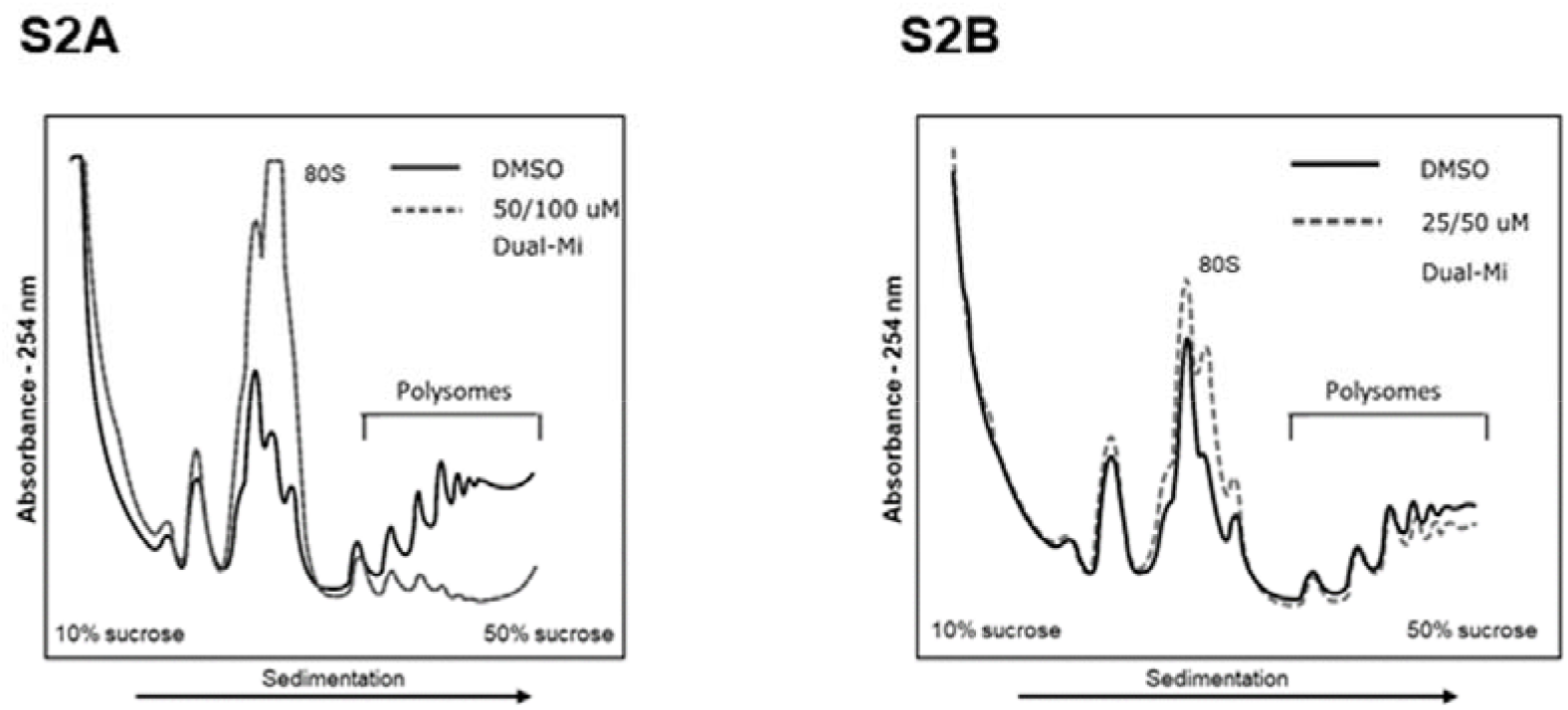
p38 and ERK1/2 dual inhibition causes polysomal collapse in MCF7 cells. **(S2A)** MCF7 cells were subjected to dual-Mi (50/100 μM p38i/ERK1/2i) or to DMSO treatment for 1-hour and polysome profiling was performed. (**S2B**) Polysome profile of the cells subjected to similar treatment but at lower concentrations of the inhibitors (25/50 μM p38i/ERK1/2i). Free ribosomal subunits (40S and 60S), monosome (80S) and the polysomes were fractionated by measuring the absorbance at 254 nm. Each graph shows treatment-curve overlaid on the vehicle control.

**Supplementary Figure S3.**
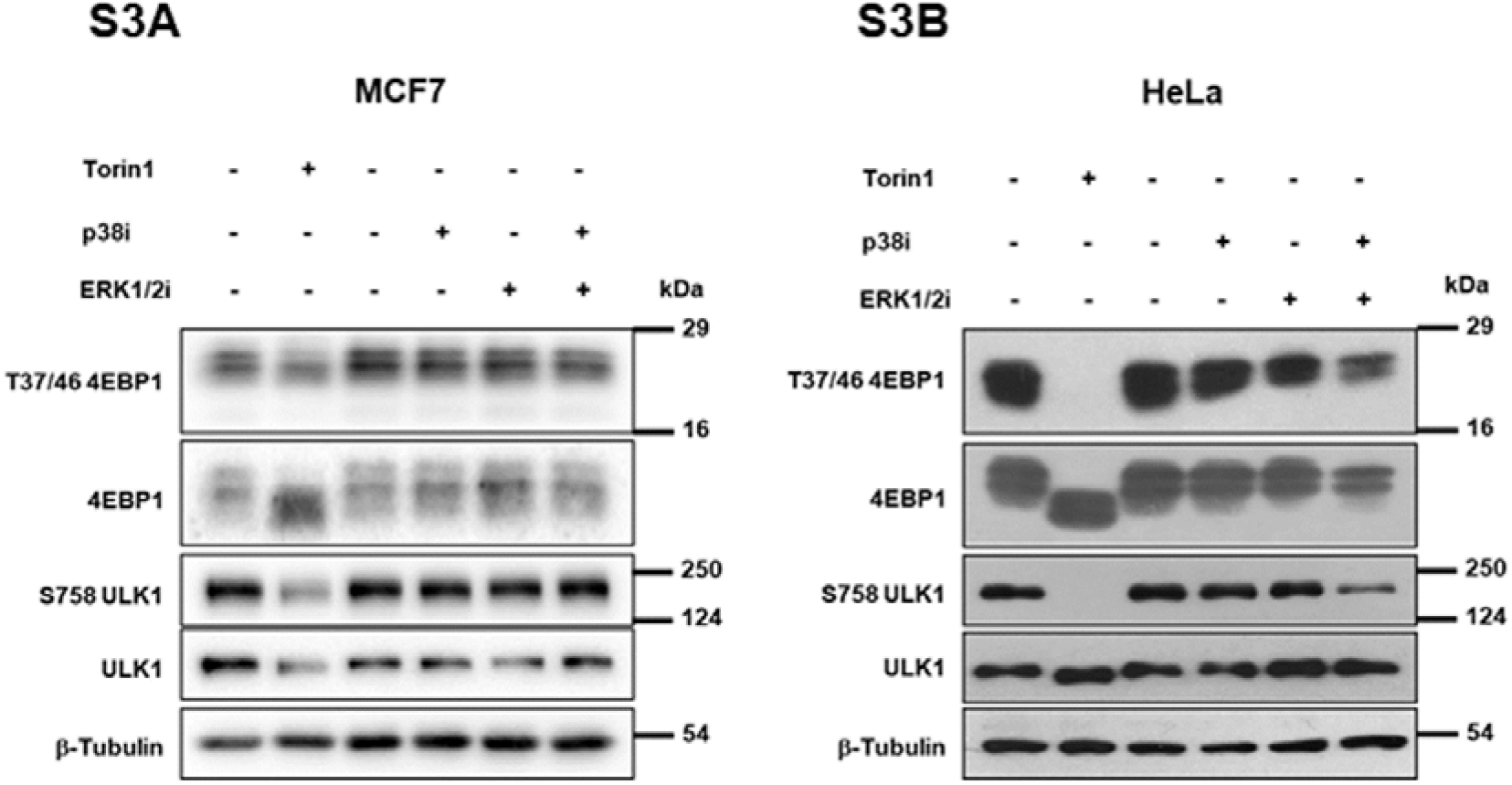
Regulation of mTORC1 pathway by p38 and ERK1/2 MAPKs is contextual. **(S3A)** Immunoblots depicting phosphorylation and expression status of 4EBP1 and ULK1 from MCF7 cells treated with DMSO vehicle control, Torin1 (750 nM), p38i (50 μM), ERK1/2i (100 μM) or Dual-Mi (50/100 μM p38i/ERK1/2i) for 1-hour. **(S3B)** Immunoblots demonstrating phosphorylation and expression status of 4EBP1 and ULK1 from HeLa cells treated with DMSO vehicle control, Torin1 (750 nM), p38i (50 μM), ERK1/2i (100 μM) or Dual-Mi (50/100 μM p38i/ERK1/2i) for 1-hour.

**Supplementary Figure S4.**
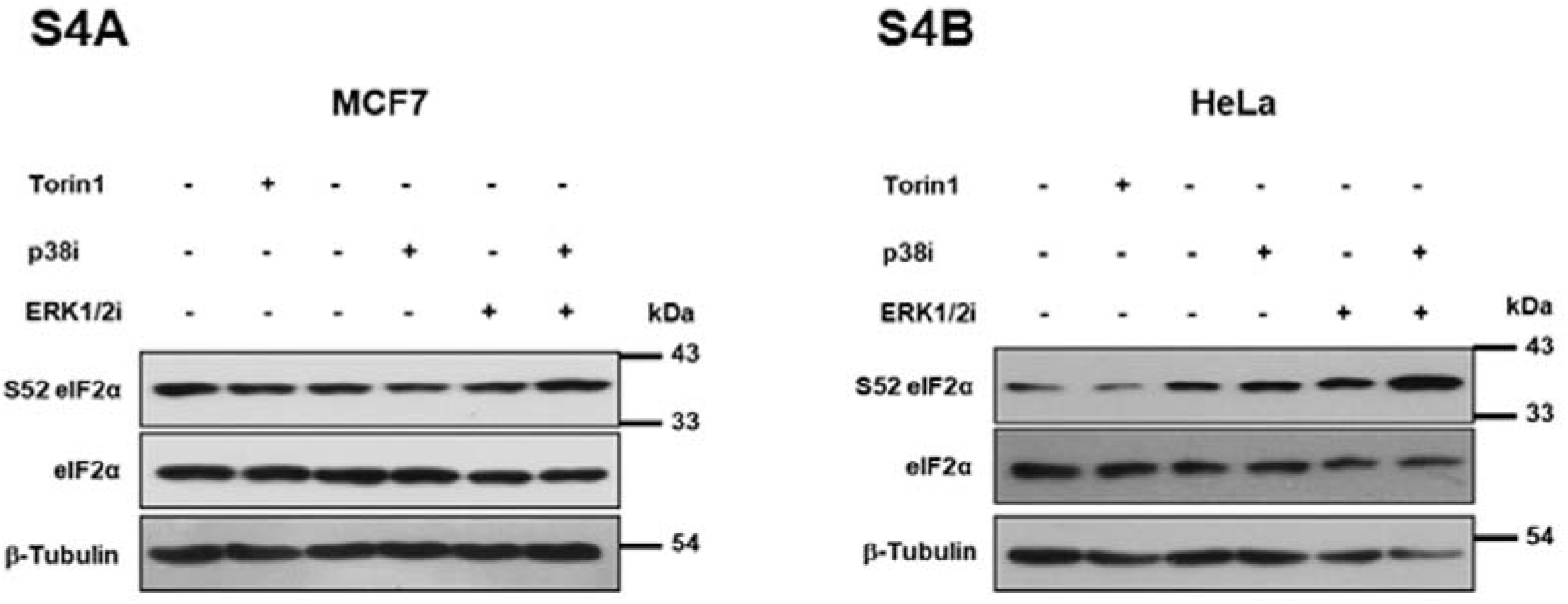
p38 and ERK1/2 dual-MAPK inhibition activates ISR signalling. **(S4A)** Immunoblots depicting phosphorylation and expression status of eIF2α from MCF7 cells treated with DMSO vehicle control, Torin1 (750 nM), p38i (50 μM), ERK1/2i (50 μM) or Dual-Mi (50/50 μM p38i/ERK1/2i) for 1-hour. **(S4B)** Immunoblots depicting phosphorylation status of eIF2α from HeLa cells treated with DMSO vehicle control, Torin1 (750 nM), p38i (50 μM), ERK1/2i (100 μM) or Dual-Mi (50/100 μM p38i/ERK1/2i) for 1-hour.

**Supplementary Figure S5.**
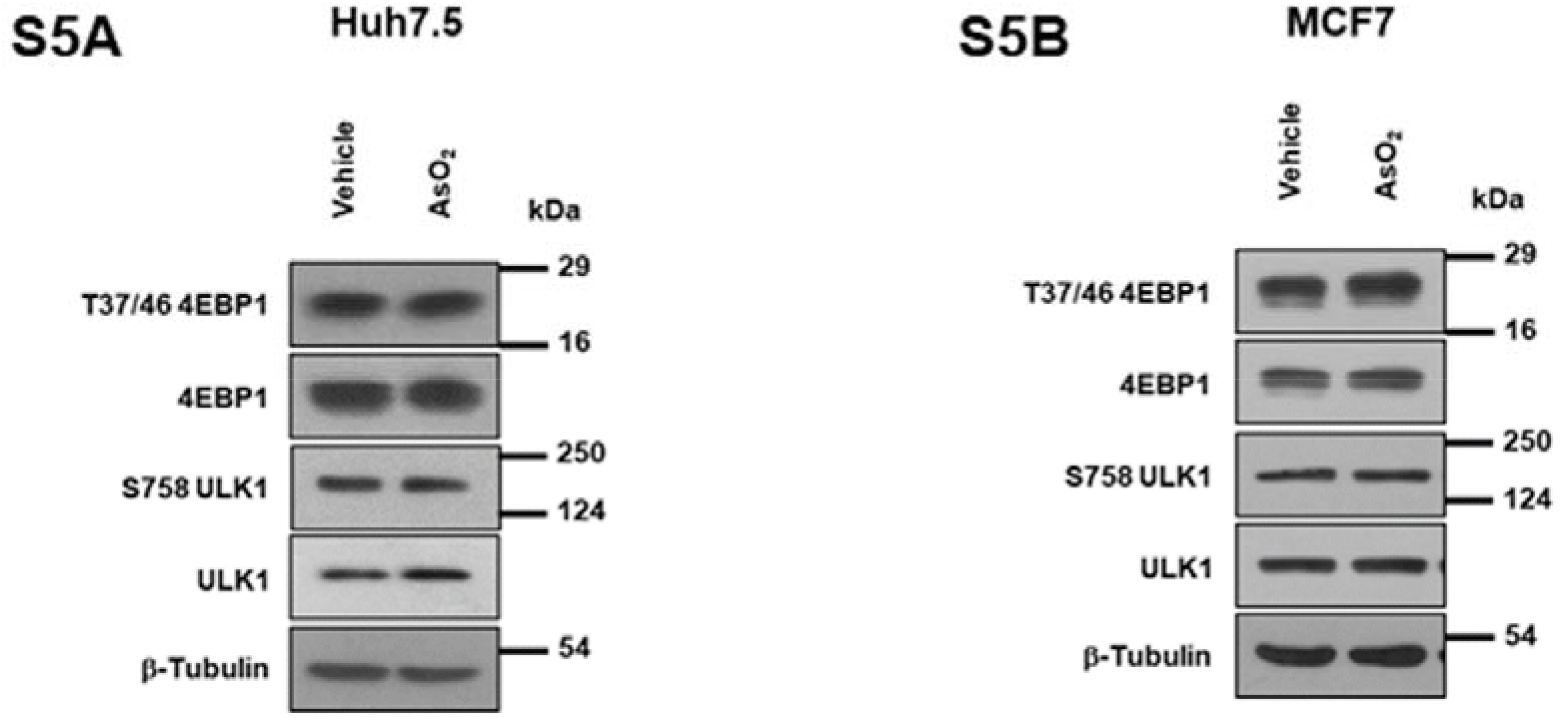
mTORC1 pathway was unaffected during sodium arsenite induced ISR. **(S5A)** Immunoblots depicting phosphorylation and expression status of 4EBP1 and ULK1 from vehicle or sodium arsenite (40 μM) treated Huh7.5 cells for 1-hour. **(S5B)** Immunoblots depicting phosphorylation and expression status of 4EBP1 and ULK1 from vehicle or sodium arsenite (40 μM) treated MCF7 cells for 1-hour.

## Notes

### Competing Interest Statement

The authors have declared no competing interest.

